# A deep learning-based iterative digital pathology annotation tool

**DOI:** 10.1101/2021.08.23.457396

**Authors:** Mustafa I. Jaber, Bing Song, Liudmila Beziaeva, Christopher W. Szeto, Patricia Spilman, Phil Yang, Patrick Soon-Shiong

**Author notes:** Corresponding author: M. I. Jaber. M. I. Jaber and B. Song contributed equally.

## Abstract

Well-annotated exemplars are an important prerequisite for supervised deep learning schemes. Unfortunately, generating these annotations is a cumbersome and laborious process, due to the large amount of time and effort needed. Here we present a deep-learning-based iterative digital pathology annotation tool that is both easy to use by pathologists and easy to integrate into machine vision systems. Our pathology image annotation tool greatly reduces annotation time from hours to a few minutes, while maintaining high fidelity with human-expert manual annotations. Here we demonstrate that our active learning tool can be used for a variety of pathology annotation tasks including masking tumor, stroma, and lymphocyte-rich regions, among others. This annotation automation system was validated on 90 unseen digital pathology images with tumor content from the CAMELYON16 database and it was found that pathologists’ gold standard masks were re-produced successfully using our tool. That is, an average of 2.7 positive selections (mouse clicks) and 8.0 negative selections (mouse clicks) were sufficient to generate tumor masks similar to pathologists’ gold standard in CAMELYON16 test WSIs. Furthermore, the developed image annotation tool has been used to build gold standard masks for hundreds of TCGA digital pathology images. This set was used to train a convolutional neural network for identification of tumor epithelium. The developed pan-cancer deep neural network was then tested on TCGA and internal data with comparable performance. The validated pathology image annotation tool described herein has the potential to be of great value in facilitating accurate, rapid pathological analysis of tumor biopsies.

## INTRODUCTION

Histopathological inspection of potentially malignant biopsies is critical across all of oncology, both in initial diagnoses (e.g. type and grade) as well as in monitoring for metastases and disease progression. Traditionally, analyses of biopsy images had been performed by expert pathologists who undergo extensive training in order to accurately identify and delineate areas of neoplasia.

Machine vision applied to digital pathology is a highly active area of research, aiming to alleviate the overwhelming demand on the short supply of experts. Digital pathology has emerged as a core technology in modern pathology that can increase accuracy of assessments while reducing delays (1). The resulting high-resolution whole-slide digital scans are ripe for application of machine vision to cell and tissue type determination.

Tissue annotations are required in order to access the power of machine vision technology and, unlike in most other machine vision tasks, pathology annotations cannot be gathered by ‘mechanical turk’ (that is, by distributing the work to many people), but rather can only be provided by the same group of experts desiring assistance. As such, there exists an explicit need to develop tools that amplify pathologists’ annotation efforts.

The first generation of annotation tools, starting with the introduction of OpenSlide (2), include a set of pixel-based digital slide annotation tools such as QuPath (3), and SlideRunner (4), Cytomine (5) and Cancer Digital Slide Archive Project (6). These systems gave the pathologist the power to view WSI, freehand outline region of interest, label, and share annotations with others. However, they lack the power of AI such as active learning that would help the annotation system evolves through continuous learning, few shot learning with would dramatically reduce the time and labor of annotating an entire WSI. Recently, OpenHI2 - open source histopathological image platform (7) announced implementing VGG, GoogLeNet, MobileNet, and Inception architectures within their system for suggestive WSI segmentation. Furthermore, the move to AI-enrich annotation tools was facilitated by the introduction of PyHIST (8); an open source WSI tissue segmentation and preprocessing command-line tool for image patch (tile) generation for machine learning applications.

Related work that uses AI includes that of Lutnick, *et al*. (9), where an iterative annotation tool for sets of WSIs has been reported. In their system, a deep learning model is trained for a certain task (semantic segmentation of WSI, for example), with a visual display of predictions provided to the pathologist for confirmation or correction after every training epoch. In Zhu *et al*. (10), a tool for human-in-the-loop deep learning of renal pathology was introduced. The human would give feedback on the optimal threshold to detect which glomeruli to retain. Their process reduced labor cost by 57% for curating large-scale target objects in WSIs. Furthermore, the concept of few shot learning was used in Medela *et al*. (11) to transfer knowledge from a well-defined source domain (colon tissue, for example) into a more generic domain composed by colon, lung, and breast tissues by using very few training histopathological images. The few shot learning approach is used as a metric learning perspective where new concepts and new representations are learned from few samples in the new domain. In Teh and Taylor’s “Learning with Less Data Via Weakly Labeled Patch Classification in Digital Pathology” (12), a reversed knowledge transfer concept was used. That is, transferable features were learned from a set of 23,916 histopathological images (arranged into 24 groups based on visual distinction). Several image patch classifiers were trained on learned features and applied on test images to suggest annotate patches with eight labels: Tumor Epithelium, Simple Stroma, Complex Stroma, Immune Cells, Debris and Mucus, Mucosal Glands, Adipose Tissue, and Background. Previously, a suggestive annotation concept was used in Yang *et al*. (13) where a deep active learning framework was also developed for biomedical image segmentation. A more recent tool, TissueWand, that is based on human-in-the-loop and histopathology image segmentation is introduced in Lindvall *et al*. (14).

An optimal annotation tool should be easy to use by pathologists and require minimal time investment. A further feature would be the ability to annotate a variety of tissue and cell types such as tumor, stroma, and lymphocyte-infiltrated regions. In addition, it should have a web-based user interface that is ideal for sharing annotations between different research groups, as they avoid the need to install specific software on multiple systems (15). To our knowledge, none of the annotation approaches described above are provided as real-time web-accessible digital pathology annotation tools that are both easy to use by pathologists and easy to integrate into machine vision systems. Herein, we describe our development and validation of a digital image annotation tool that meets the criteria described above and demonstrate that it accurately reproduces the gold standard masks generated by pathologists. Our proposed image annotation tool aims to close the translation gap of artificial intelligence (AI) applications in digital pathology by increasing pathologist awareness of AI strengths and weaknesses in the domain of image annotation (16). Our proposed image annotation depends on the pathologist in the iterative process and in the review of generated masks and its interpretation (17). The proposed tool greatly reduces annotation time from hours to a few minutes, while maintaining high fidelity with human-expert manual annotations. Our active learning-based tool can be used for a variety of pathology annotation tasks including masking tumor, stroma, and lymphocyte-rich regions, among others.

## METHODS

### Nant’s digital pathology annotation tool

Figure 1 shows that the developed systems has several components that may run on one or multiple computing machines. A whole side image (WSI) is generated via a digital slide scanner or uploaded to the annotation system via a web server. The image is saved on the file server. Upon receiving the image file, the web server estimates tissue regions (using global thresholding) and cuts them into non-overlapping patches of size 100 square micron (typically > 10K patches). The web server also initiates one or more GPUs to compute patch descriptors (1D-logit vectors in ImageNet-based ResNet34). When ready, the pathologist (user) is asked to select several positive (and negative) examples; say, several tumor and non-tumor regions. A digital classifier takes pathologist’s input to classify all 1D descriptors of tissue patches into positive (tumor) and negative (non-tumor) regions. Generated makes are then shown overlaying input WSI. Pathologist is instructed to examine the generated binary mask and to select more positive / negative patch samples if needed to refine output mask. In our implementation, output mask refinement is done in less than 30 seconds for most WSIs. Several iterations of mask refinement are usually required to generate a satisfactory final mask. A block diagram of the developed annotation algorithm is shown in Figure 2.

**Fig. 1.**
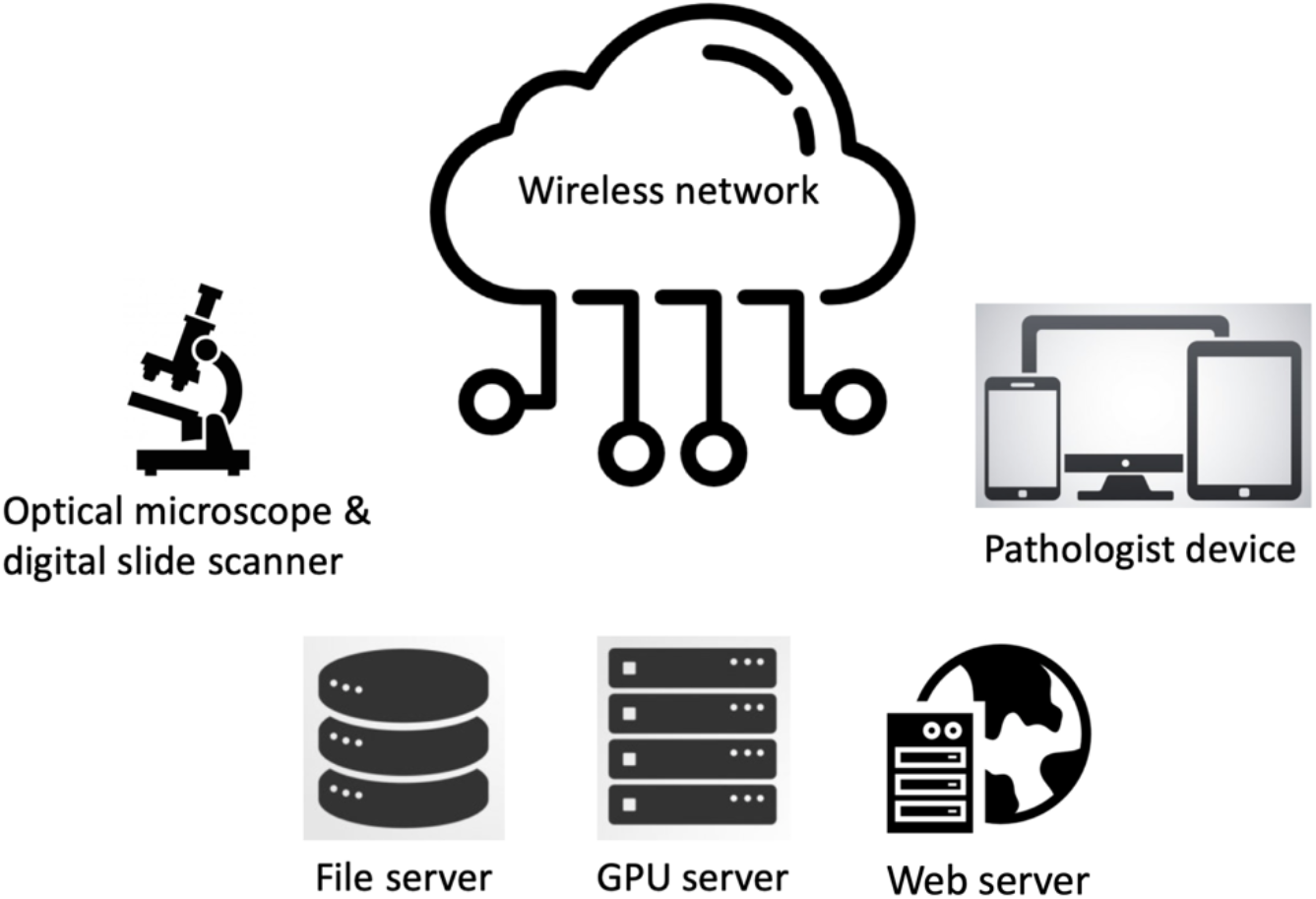
An overview of the proposed digital pathology annotation system.

**Fig. 2.**
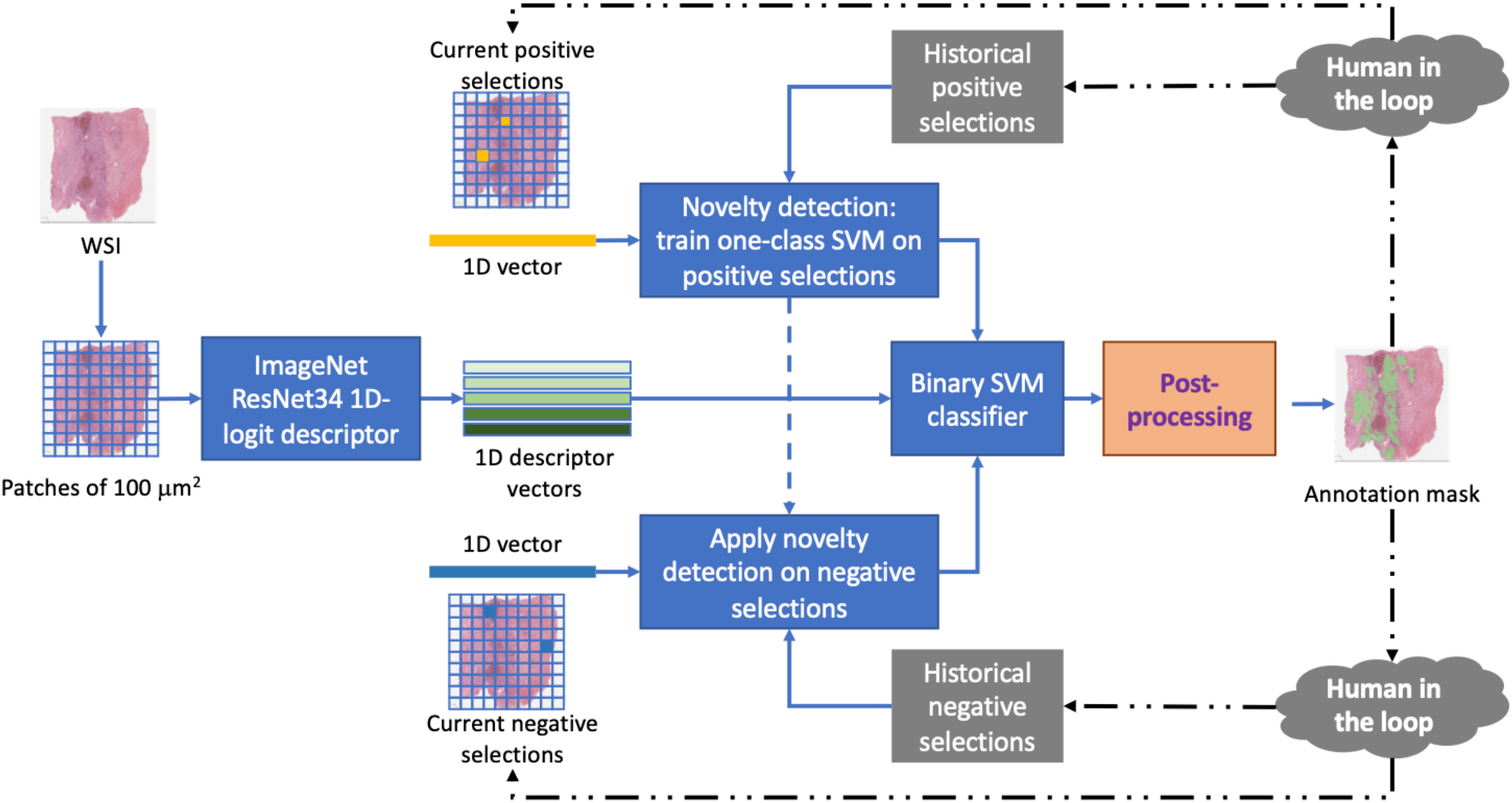
A block diagram of the annotation algorithm. Preprocessing

The preprocessing module identifies regions of interest (tissue areas) in input whole slide images (WSIs) and cuts them into non-overlapping patches, which are then described as ID vectors using a deep learning pre-trained convolutional neural network.

### Tissue mask

Segmentation of input WSIs into foreground and background regions is the first preprocessing step applied to an input image. The WSI is first converted from RGB to gray-scale color space. It is then scaled and cashed using at a low-resolution (low magnification) level. The image is then low-pass filtered (Gaussian average filter) before binarization. Otsu (18, 19), Yen (20), and global thresholding (21) methods are all applied to the input image at low-resolution. These three thresholding methods are empirically combined to generate a tissue mask.

### Valid patch queue

Several computational threads are initiated to crop valid tissues into non-overlapping square patches of size 100 micron. These image patches are sent to the GPUs for further processing.

### Patch descriptor

A 1D image descriptor is extracted per patch using a pre-trained ImageNet ResNet34 module. Vector values in Logit layer, that is, the vector of raw (non-normalized) predictions that the pre-trained ResNet34 model generates, are used to transform 2D color patches into 1D vectors.

### Novelty detection: one-class SVMs

At this stage, the WSI is ready for annotation. The system aims to generate a robust binary mask of positive and negative regions based on minimal positive and negative selections (mouse clicks) from the user (pathologist). In order to achieve this, the system utilizes all historical patch (positive and negative) selections to progressively chooses more discriminative patches that contribute to better model performance. Note that our samples include current positive selections, current negative selections, historical positive selections, and historical negative selections.

A novelty detection - unsupervised learning mechanism - module based on one-class SVM is applied on positive samples to prune historical positive selections that are similar to current positive selections. The same novelty detector is also used to remove historical negative selections that are similar to the current positive selections. Organized positive and negative training sets are used to power a binary SVM classifier on current WSI data.

Figure 3A shows examples (in 2D space) of application of the novelty detection module to create positive training set for the binary SVM classifier. In the example, we see training one-class SVMs on current positive selections generates one (or more) “learned” frontiers. Such frontiers are used as boundaries to recruit similar descriptors from historical selections. Note that the 1D descriptor in this work is 512, which is the Logit layer of ResNet34 model.

**Fig. 3.**
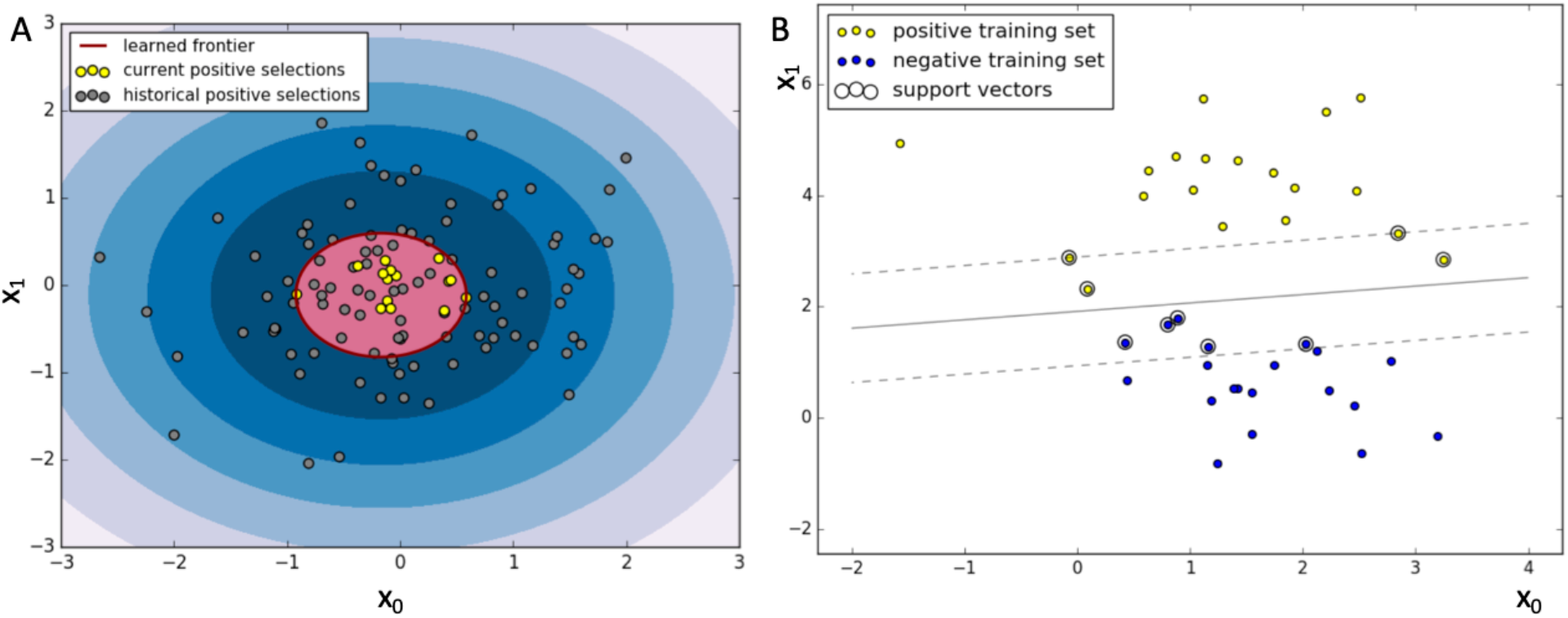
An illustrated example of novelty detection and binary linear SVM classifier. (A) Novelty detection trained on 15 current positive selections pruned 26 (of 100) historical positive selections to yield a positive training set (for binary SVM classifier). (B) Binary linear SVM classifier on positive and negative training sets. [Code to generate Fig. 3 is adopted from https://scikit-learn.org]

### Binary SVM classifier

The positive training set (for binary SVM classifier) includes all current positive selections and similar ones from historical positive selections while the negative training set (for binary SVM classifier) includes all current negative selections and historical negative selections (except similar ones to current positive selections, if any). With enough samples in the training set, a robust binary linear SVM classifier is trained. Figure 3B illustrates an example of a two-class SVM classifier where negative support vectors (circled samples that cross the bottom dashed line) and positive support vectors (circled samples that cross the top dashed line) maximize the margin width (the width between the two parallel dashed lines). The hyperplane is the solid line shown in Figure 3B at the midpoint between the two dashed lines.

The trained two-class SVM described can then be used to classify test patches as either being a positive patch (positive for cancer, for example) if the patch is on the positive side of the hyperplane, or a negative patch (negative for cancer, for example) if the patch is on the negative side of the hyperplane.

### Post-processing

The binary classification step generates a patch-level mask. To further improve the generated mask, an optional post-processing step for grouping can be developed. That is, test patches that have been determined to be positive can be grouped with other neighboring positive patches. Similarly, the test patches that have been determined to be negative can be grouped with other neighboring negative patches. From such groupings, convex hulls can be generated across adjacent patches, and used to detect and correct locally mis-classified patches within said convex hulls. This post-processing step can enhance annotation masks when large blobs are expected as in tumor annotations. However, it is not of use and should not be applied in case of lymphocyte annotations as they are typically scattered and / or isolated patches.

### Preloaded database for tumor epithelium detection

The developed system is a generic patch-level annotation tool for WSIs. It can be used to annotate tumor epithelium, lymphocyte regions, stroma tissue, and more. In the case of tumor annotation, a new user may choose to use the system with no historical selections or opt for a preloaded database of positive and negative selections as shown in Figure 4.

**Fig. 4.**
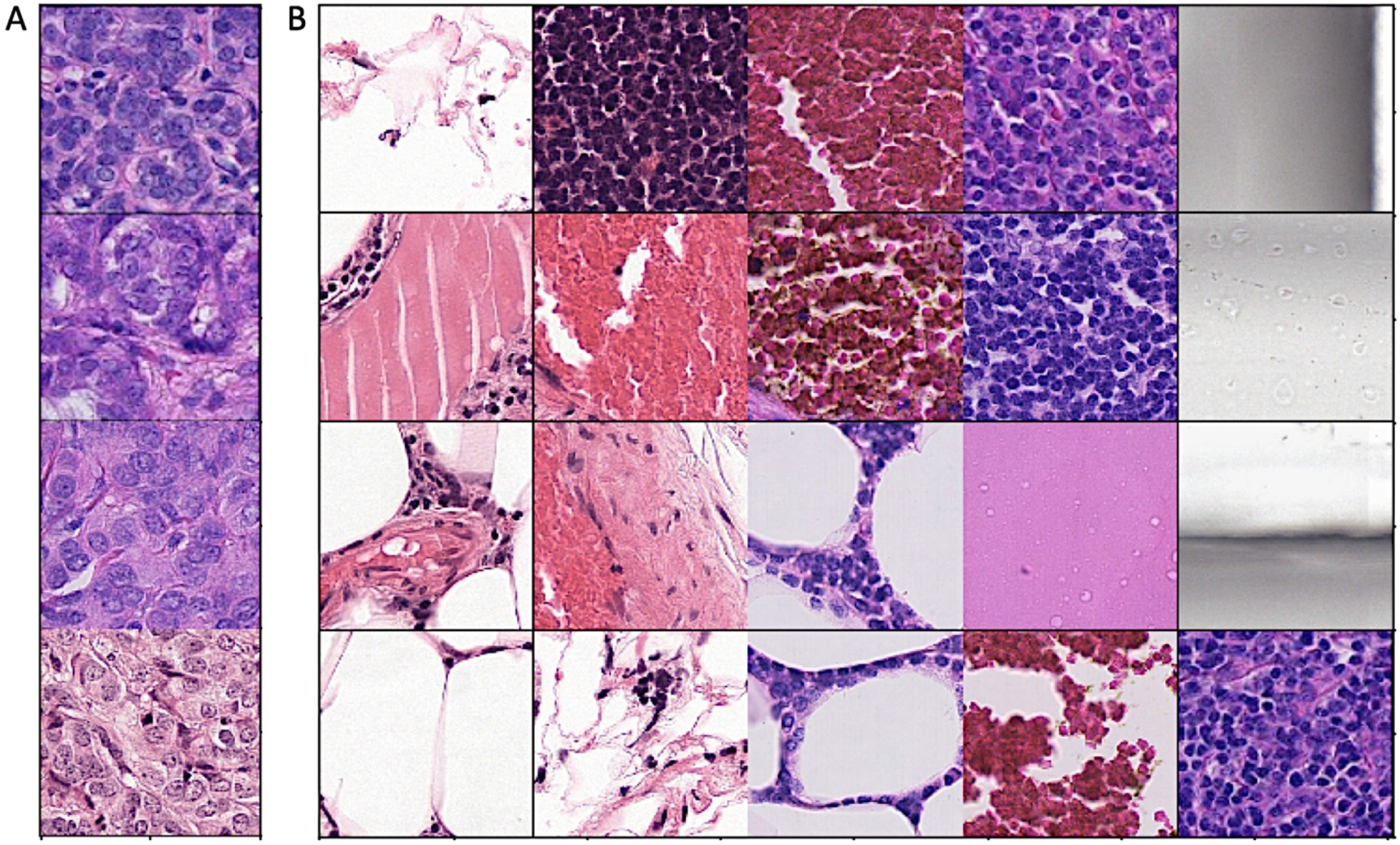
Pre-loaded historical data sets. (A) Preloaded positive database. (B) Preloaded negative database.

Using this custom preloaded database minimizes the number of initial negative selections to cover background, scanning defects, and no-cell areas. It also minimizes the number of positive selections as four tumor patches are used as examples.

Note that all new (positive and negative) selections by the pathologist are saved and used as “historical selections” for future image annotations. This active learning scheme would help the annotation system evolves through continuous learning.

### Annotation Tool Performance

Our goal was to develop an annotation tool that would yield similar (if not exact) annotation masks to those generated manually by a pathologist using other tools. In order to evaluate how our tool compares to pathologist annotation, a simulation experiment was run using the CAMELYON16 database where positive (tumor) and negative (non-tumor) selections were randomly chosen per WSI to achieve a desired F1 score.

### Gold standard tumor masks

The CAMELYON16 database provided freehand tumor annotations of 90 tumor-containing WSIs. In our experiment, we converted these annotations to patch-based ones using 100 square micron patches. Where annotations of tumor/non-tumor regions were provided as free-hand curves, these divide some patches into tumor and non-tumor portions. In such cases, patches were included in the tumor mask if they contained at least 40% tumor content to retain as much tumor area as possible. The average number of tissue patches (of size 100 square micron) in these 90 WSIs is 17512.44 with 829.12 average patches identified as tumor in the CAMELYON16 gold standard masks. That is an average of 4.73% tumor content.

### Experiment setup

The goal of the simulation experiment was to quantify the minimum number of positive and negative selections needed to achieve a desired tumor mask quality using the developed annotation tool. Mask quality is represented here by F1 score. The F1 score is generated by comparison of the developed mask to the patch-based gold standard tumor mask. In the experiment, we started with the preloaded positive and negative historical selections shown in Figure 4, and incrementally increased the number of positive and negative selections to achieve desired F1 scores as shown in Table 1. In this simulation, we placed the positive selection(s) within the given patch-based gold standard tumor area at random. Similarly, negative selections are placed outside tumor area randomly. To ensure robust performance, selections sets (positive and negative) were repeated 20 times. That is, if the simulation was testing masks generated using one positive and one negative selection, these selections were randomly placed 20 times and the generated masks were evaluated each time. We anticipate that if our annotation system is used by a pathologist, he/she will use educated guesses based on previous experience to place selections. However, this is not possible in our simulation and thus repetition was used.

**Table 1:** Annotation tool performance evaluation

| F1 score | >0.60 | >0.65 | >0.70 | >0.75 | >0.80 | >0.85 | >0.90 |
| --- | --- | --- | --- | --- | --- | --- | --- |
| Tumor selections | 1.54 | 1.82 | 2.18 | 2.73 | 3.77 | 4.20 | 5.00 |
| Nontumor selections | 2.86 | 4.66 | 6.60 | 8.00 | 12.33 | 13.39 | 17.90 |
| % out of 90 WSIs | 100.0 | 100.0 | 100.0 | 98.89 | 95.56 | 87.78 | 75.56 |

### Performance evaluation results

Our simulations showed that one positive selection (i.e. one mouse click) was adequate to generate a tumor mask with F1 scores > 0.75 in 40/90 (44.4%) tumor WSIs, using just the preloaded historical positive and negative patches for tumor epithelium detection and no negative selections. By making a total of six selections (e.g. two positive and four negative), F1 scores of > 0.75 were found for an increased number of WSIs, to 55/90 (61.11%) in the test WSIs. Table 1 shows that an average of 2.73 positive selections and 8.00 negative selections were sufficient to generate tumor masks with acceptable quality (F1 score > 0.75) in 89 out of 90 CAMELYON16 tumor WSIs used in this experiment. The other image achieved F1 score of 0.71 with 25 tumor and 125 nontumor selections.

Figure 5 shows two gold standard tumor masks of CAMELYON16 WSIs that were reproduced using our developed annotation tool with just one positive and one negative patch selection (i.e. two mouse clicks). Note that although the WSIs contains at least four distinct tumor regions, one positive selection in one of the tumor regions was adequate to capture the majority of tumor area across the WSI.

**Fig. 5.**
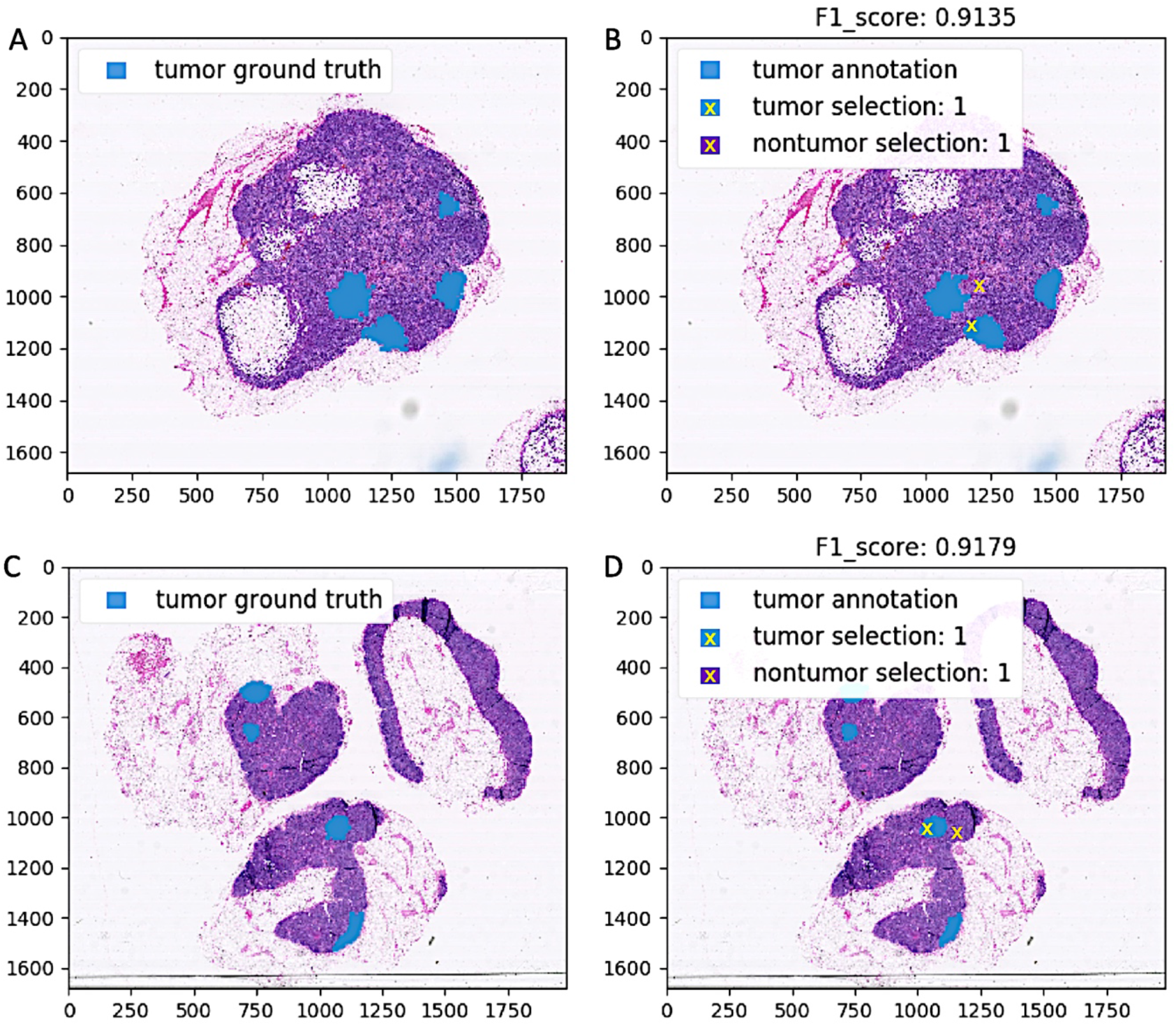
Re-produced gold standard tumor masks of two CAMELYON16 WSIs. Two gold standard tumor masks (A, C) were re-produced (B, D) using the developed annotation tool with one positive (yellow x) and one negative (orange x) patch selection.

### Use case: Identification of tumor epithelium

Cancerous cells are mostly found in the epithelium area in many cancer types. Detection of epithelial tumor regions in standard Hematoxylin and Eosin stained histology images is an essential step. This could be used for the analysis of tissue architecture (22), or to predict overall survival and outcome (23), among other applications. When considering pan-cancer image-based epithelial tumor detection tool, this problem becomes challenging due to the high variability in chromatin texture of the tumor nuclei, in addition to their irregular sizes and shapes (22). Here we explored whether large deep learning framework such as inception network are capable of handling such complexities.

### Developed Solution

The human-in-the-loop annotation tool presented in Section II was used to develop patch-level tumor masks for 433 WSIs from ten different TCGA cohorts. This set was further used to train an inception v3 network as shown in Figure 6.

**Fig. 6.**
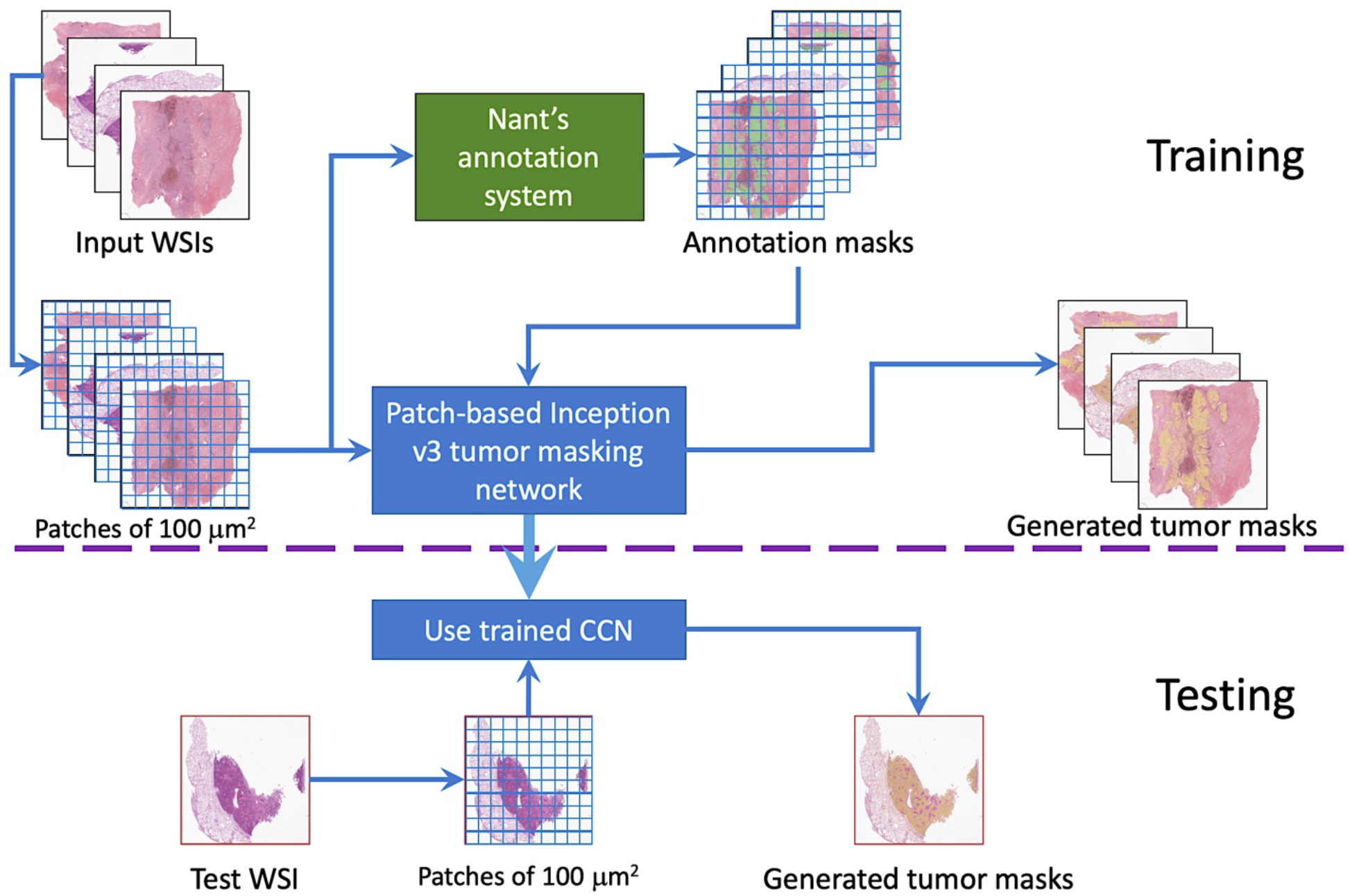
Block diagram of tumor epithelium identification.

Stochastic gradient descent in PyTorch with momentum of 0.9 and weight decay of 1e-4 were used to minimize the training loss. An exponential moving average parameter of 0.85 and learning rate of 1.375e-05 were also used. Equal numbers of tumor (positive) and nontumor (negative) patches were randomly sampled from training WSIs to give 443,392 patches (443,392 = 512 patch x 433 WSI x 2 class). Our data augmentation strategy was to randomly rotate training patches by 0°, 90°, 180°, or 270° whenever they are used in a new training epoch. The inception v3 trained system gave the highest patch-level classification accuracy for the 18 WSI validation set after 4,443,920 steps; that is, with ten epochs and 10K steps.

### Identification Network

The trained CNN was used to generate tumor masks automatically for three test sets. The first contained 82 TCGA WSIs from the same (ten) cohorts used in the training set, the second contained 36 TCGA WSIs from eight cohorts not used in the training set, and the last one contained 186 clinical WSIs obtained from the Nant database with true positive tumor regions annotated by a pathologist.

### Identification of Tumor Epithelium on TCGA Data

Samples form ten TCGA cancer cohorts were used to train our deep net for tumor identification. Other (not used in training) diagnostic WSI samples from these ten TCGA cohorts were used to evaluate its classification performance. Per-cancer accuracy is shown in Figure 7A. The average number of patches (square patches of size 100 micron) in this 82 test WSI set is 22,023 per WSI. TCGA-COAD colon cancer WSIs have the least number of average patches per WSI with 15,327 while TCGA-BRCA breast cancer has the largest WSI size with 27,448 patches per WSI on average. Figure 7A shows the trained deep net performed well on TCGA-BLCA Bladder and TCGA-BRCA breast cancers while improvement is needed for TCGA-PAAD pancreas cancer samples.

**Fig. 7.**
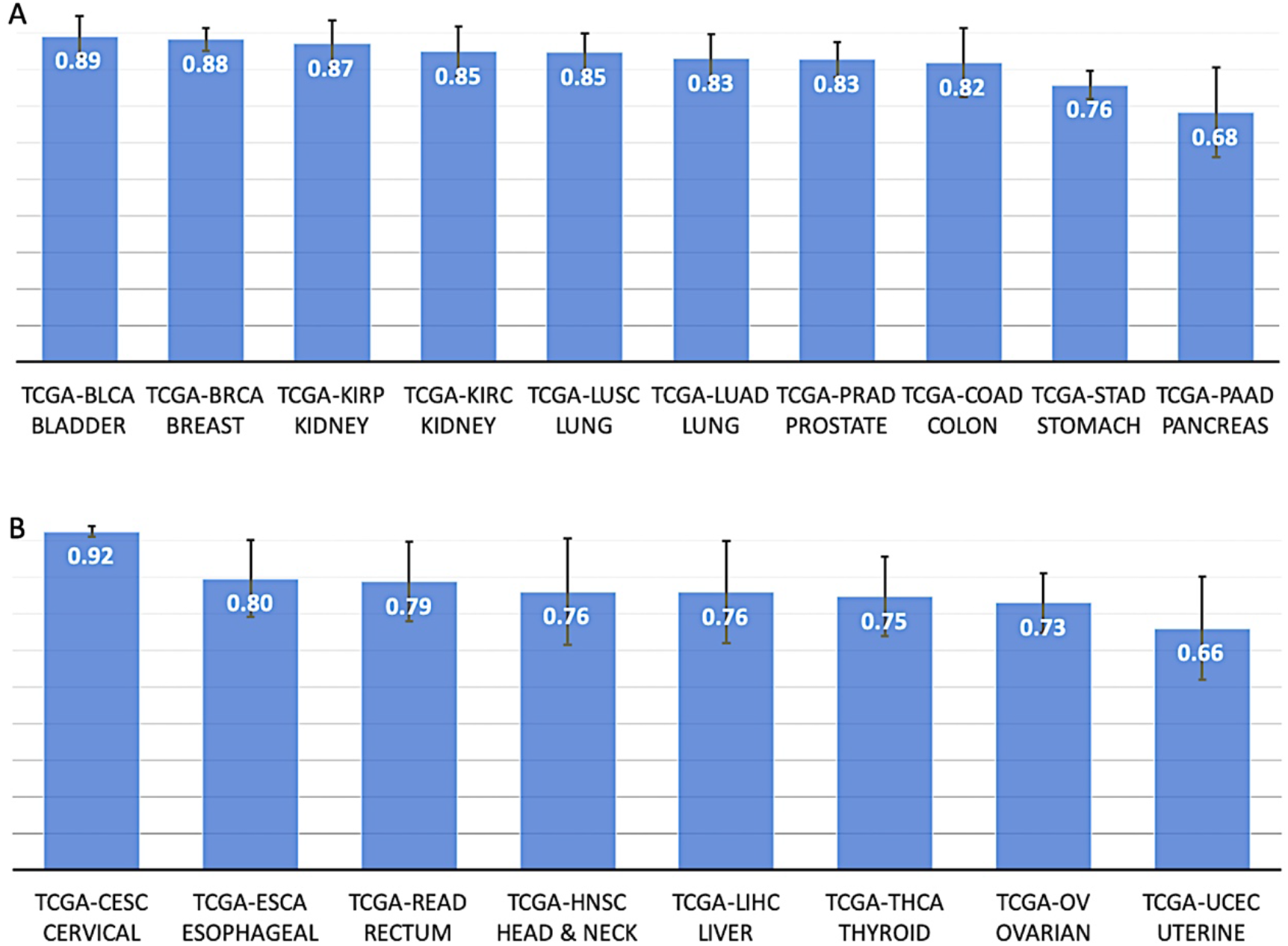
Accuracy of identification of tumor epithelium performance. (A) Cancer types in training data are shown. (B) Cancer types not in training data are shown, The accuracy of tumor epithelium identification based on testing a set of 118 TCGA WSIs from different cancer cohorts is high, matching deep-net-generated tumor masks to pathologists annotations.

We further demonstrate that the trained tumor-detecting network can generalize to unseen samples by evaluating diagnostic WSIs of cancers not included in its training data (Figure 7B). The best classification accuracy was achieved in TCGA-CESC cervical cancer samples while TCGA-UCEC uterine cancer samples need improvement. The average number of square patches in this 36 WSI test set was 21,673 with 12,914 patches (on average) in TCGA-READ rectum cancer WSIs and 34,855 patches in TCGA-UCEC uterine WSIs, respectively.

Figure 8 shows visual results of selected test WSIs from TCGA datasets with total patch count and accuracy. The green mask represents human-in-the-loop annotations while the yellow mask is generated via the trained deep net. Patches of size 750 × 366 pixels at 5X magnification are shown in the figure.

**Fig. 8.**
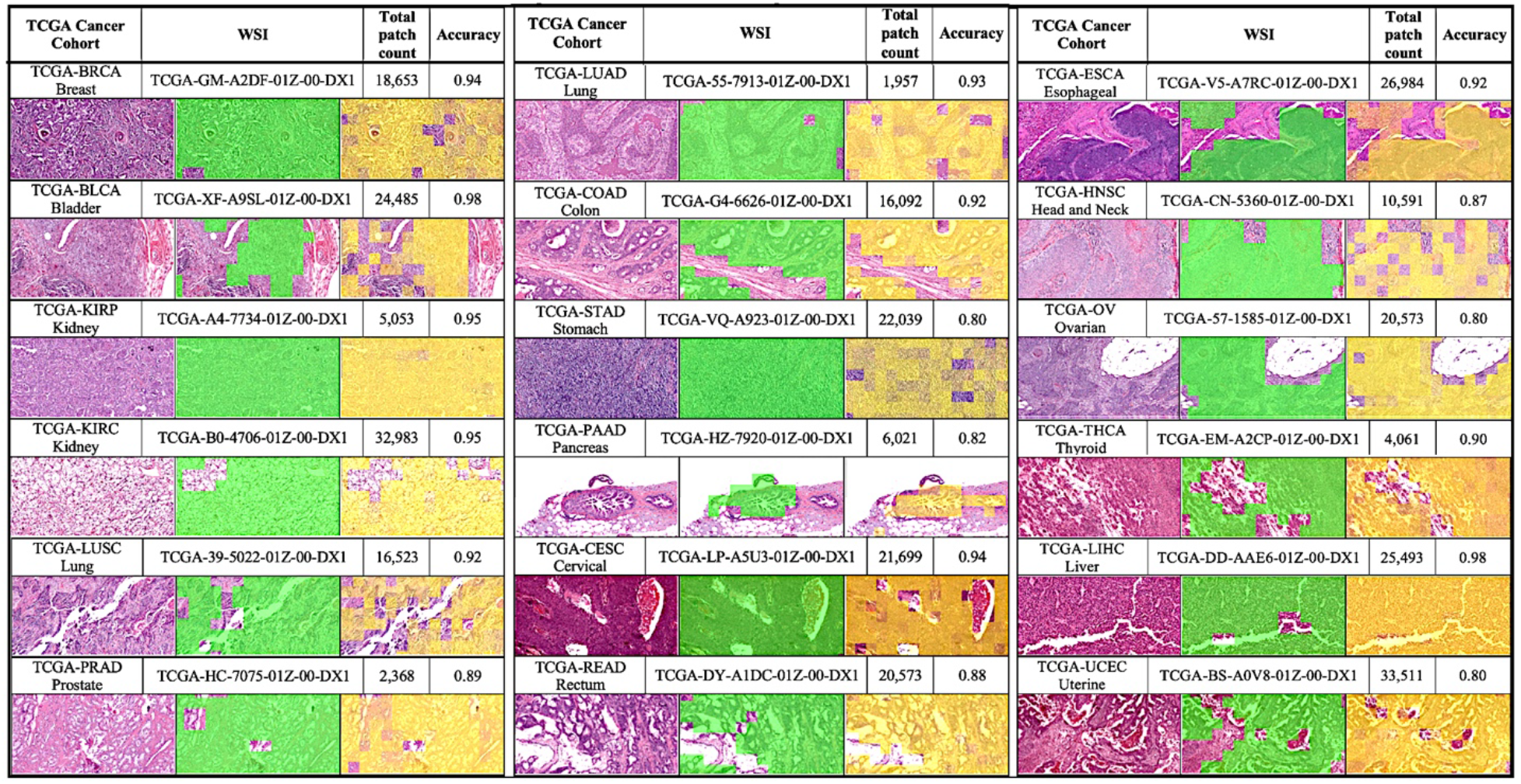
Identification of tumor epithelium on visual examples from our TCGA testing set with different cancers. Selected regions of tumor, pathologist’s annotations using Nant’s annotation system (green masks), and deep-net-generated tumor masks (in yellow) are also shown at 5X magnification.

### Identification of Tumor on internal Data

In clinical samples obtained from Nant, sufficient tumor-rich area (>80% tumor) was marked by pathologists for -omic characterization at a resolution suitable for laser-capture microdissection (LCM). As such, if sufficient tumor tissue had been marked the pathologist could stop, thus regions outside of annotations are not necessarily negative for tumor. To overcome this limitation, we used true positive tumor as a performance criterion. That is, the deep net (trained on TCGA data only) is expected to identify all regions (in Nant’s internal data) outlined by the pathologist as tumor. Per-cancer tumor identification results on Nant’s database (186 clinical WSIs) are shown in Figure 9 while two visual examples on lung and esophageal tissues are shown in Figure 10.

**Fig. 9.**
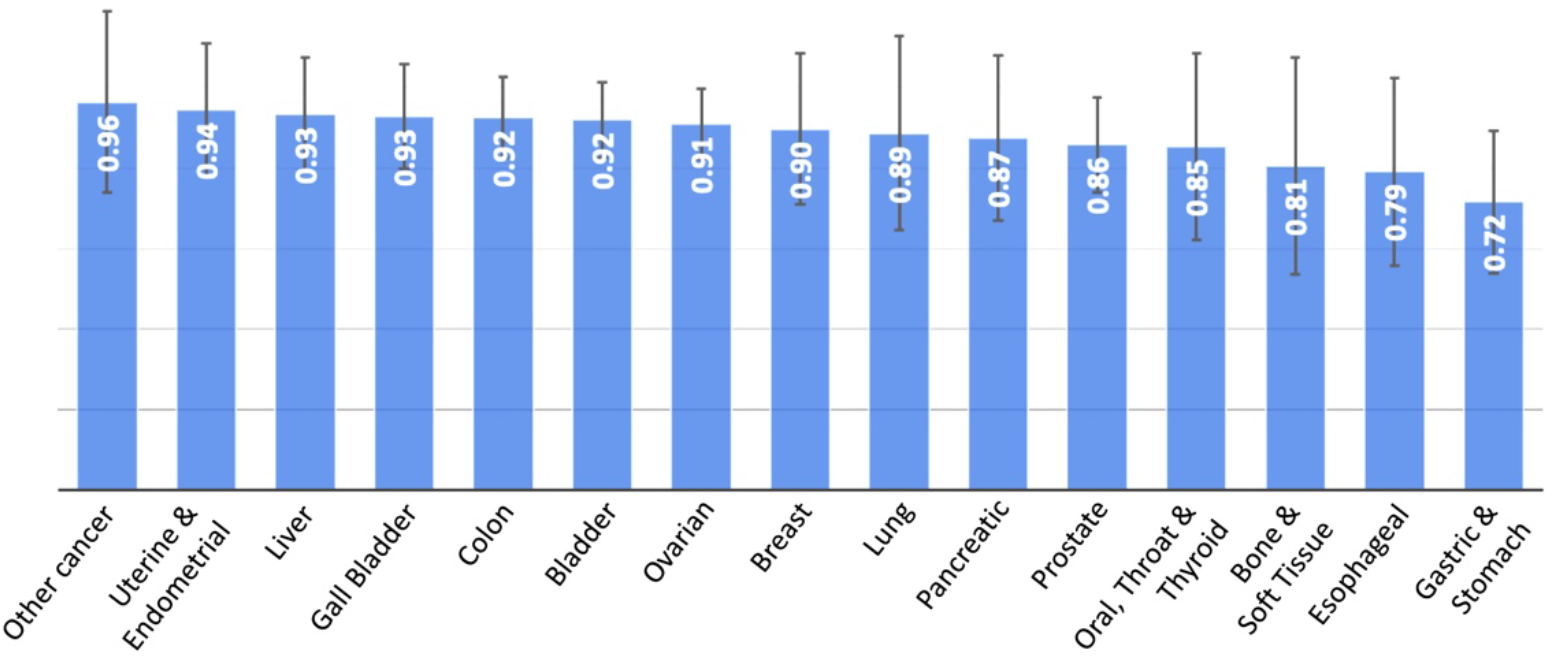
Identification of tumor performance on Nant’s testing set of 186 WSIs from different cancers. True positive tumor regions that match pathologist’s annotations are used as a performance metric here.

**Fig. 10.**
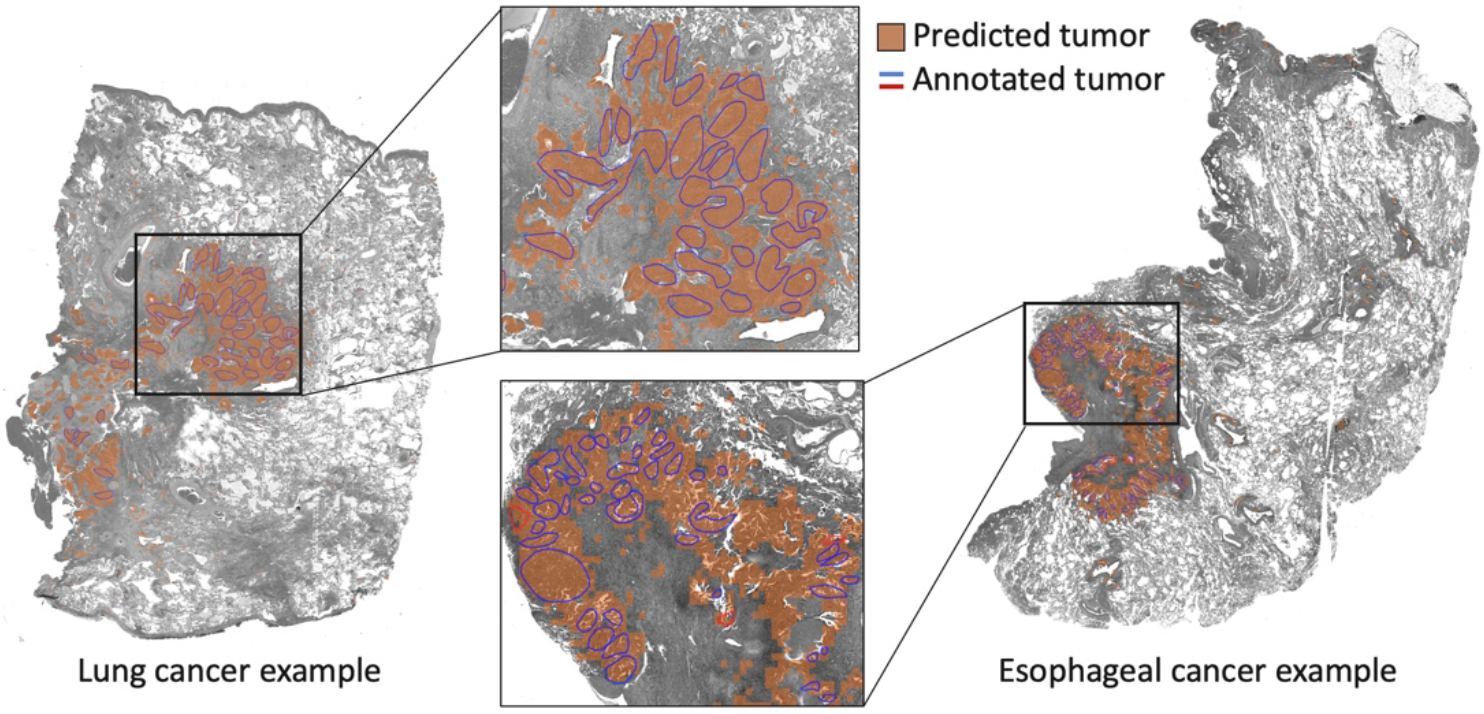
Identification of tumor performance on Nant’s WSIs shows a high true positive rate between deep-net-generated tumor masks and pathologist’s annotations.

## CONCLUSION

Well-annotated exemplars are an important prerequisite for supervised deep learning schemes. Unfortunately, generating these annotations is a cumbersome and laborious process, due to the large amount of time and effort needed (24). We present a deep-learning based iterative digital pathology annotation tool that is both easy to use by the pathologist and easy to integrate with patch-based deep learning systems. Our pathology image annotation tool greatly reduces the time needed to generate patch-level masks from hours to a few minutes while maintaining mask / annotation quality. The tool has the option to continuously store pathologists’ selections to help guide performance in future WSIs as an active learning framework. Our tool can be used to annotate tumor, stroma, and lymphocyte regions, among others.

## ACKNOWLEDGMENT

The authors would like to thank Clive R. Taylor, M.D., DPhil; and John Yahya Elshimali, M.D., FCAP, FASCP for their digital pathology support.

## REFERENCES

1. Niazi MKK, Parwani AV, Gurcan MN. Digital pathology and artificial intelligence. The Lancet Oncology. 2019;20(5):e253–e61.

2. Goode A, Gilbert B, Harkes J, Jukic D, Satyanarayanan M. OpenSlide: A vendor-neutral software foundation for digital pathology. Journal of pathology informatics. 2013;4:27.

3. Bankhead P, Loughrey MB, Fernández JA, Dombrowski Y, McArt DG, Dunne PD, et al. QuPath: Open source software for digital pathology image analysis. Sci Rep. 2017;7(1):16878.

4. M. A, C. B, R. K, A. M. SlideRunner. In: A. M, T. D, H. H, K. M-H, C. P, T. T, editors. Bildverarbeitung für die Medizin 2018 Informatik aktuell. Berlin, Heidelberg: Springer Vieweg; 2018. p. 309–14.

5. Marée R, Rollus L, Stévens B, Hoyoux R, Louppe G, Vandaele R, et al. Collaborative analysis of multi-gigapixel imaging data using Cytomine. Bioinformatics. 2016;32(9):1395–401.

6. Gutman DA, Cobb J, Somanna D, Park Y, Wang F, Kurc T, et al. Cancer Digital Slide Archive: an informatics resource to support integrated in silico analysis of TCGA pathology data. J Am Med Inform Assoc. 2013;20(6):1091–8.

7. Puttapirat P, Zhang H, Deng J, Dong Y, Shi J, He H, et al., editors. OpenHI2 — Open source histopathological image platform. 2019 IEEE International Conference on Bioinformatics and Biomedicine (BIBM); 2019 18–21 Nov. 2019.

8. Muñoz-Aguirre M, Ntasis VF, Rojas S, Guigó R. PyHIST: A Histological Image Segmentation Tool. PLoS Comput Biol. 2020;16(10):e1008349.

9. Lutnick B, Ginley B, Govind D, McGarry SD, LaViolette PS, Yacoub R, et al. An integrated iterative annotation technique for easing neural network training in medical image analysis. Nat Mach Intell. 2019;1(2):112–9.

10. Z Z, Y L, R D, H Y, B FA, Y H. EasierPath: An Open-Source Tool for Human-in-the-Loop Deep Learning of Renal Pathology. In: Cardoso J, editor. Interpretable and Annotation-Efficient Learning for Medical Image Computing. 12446: Springer Chem; 2020.

11. Medela A, Picon A, Saratxaga CL, Belar O, Cabezón V, Cicchi R, et al., editors. Few Shot Learning in Histopathological Images:Reducing the Need of Labeled Data on Biological Datasets. 2019 IEEE 16th International Symposium on Biomedical Imaging (ISBI 2019); 2019 8–11 April 2019.

12. Teh EW, Taylor GW, editors. Learning with Less Data Via Weakly Labeled Patch Classification in Digital Pathology. 2020 IEEE 17th International Symposium on Biomedical Imaging (ISBI); 2020 3–7 April 2020.

13. Yang L, Zhang Y, Chen J, Zhang S, Chen D, editors. Suggestive Annotation: A Deep Active Learning Framework for Biomedical Image Segmentation. MICCAI; 2017.

14. Lindvall M, Sanner A, Petré F, Lindman K, Treanor D, Lundström C, et al. TissueWand, a Rapid Histopathology Annotation Tool. Journal of pathology informatics. 2020;11:27.

15. Abels E, Pantanowitz L, Aeffner F, Zarella MD, van der Laak J, Bui MM, et al. Computational pathology definitions, best practices, and recommendations for regulatory guidance: a white paper from the Digital Pathology Association. J Pathol. 2019;249(3):286–94.

16. Steiner DF, Chen PC, Mermel CH. Closing the translation gap: AI applications in digital pathology. Biochim Biophys Acta Rev Cancer. 2020;1875(1):188452.

17. Aeffner F, Zarella MD, Buchbinder N, Bui MM, Goodman MR, Hartman DJ, et al. Introduction to Digital Image Analysis in Whole-slide Imaging: A White Paper from the Digital Pathology Association. Journal of pathology informatics. 2019;10:9.

18. Vala HJ, editor A Review on Otsu Image Segmentation Algorithm Miss 2013.

19. Otsu N. A Threshold Selection Method from Gray-Level Histograms. IEEE Transactions on Systems, Man, and Cybernetics. 1979;9(1):62–6.

20. Yen JC, Chang FJ, Chang S. A new criterion for automatic multilevel thresholding. IEEE Trans Image Process. 1995;4(3):370–8.

21. Chaubey AK, editor Comparison of The Local and Global Thresholding Methods in Image Segmentation 2016.

22. Sirinukunwattana K, Raza SEA, Tsang Y-W, Snead D, Cree I, Rajpoot N. A Spatially Constrained Deep Learning Framework for Detection of Epithelial Tumor Nuclei in Cancer Histology Images. Wu G, Coupé P, Zhan Y, Munsell B, Rueckert D, editors: Springer Chem; 2015.

23. Alom MZ, Aspiras T, Taha TM, Asari VK, Bowen TJ, Billiter D, et al. Advanced Deep Convolutional Neural Network Approaches for Digital Pathology Image Analysis: a comprehensive evaluation with different use cases. arXivorg. 2019.

24. Tao Y, Peng Z, Krishnan A, Zhou XS. Robust Learning-Based Parsing and Annotation of Medical Radiographs. IEEE Transactions on Medical Imaging. 2011;30(2):338–50.

