## SUPPLEMENTARY Figure S1 and Figure S2 for "A deep learning-based iterative digital pathology annotation tool"

**FOR**

### SUPPLEMENTARY Figure S1

Figure S1 shows the one image example that our annotation tool failed to re-generate pathologist gold standard with minimum number of selections. This image achieved F1 score of 0.71 with 25 tumor and 125 nontumor selections.

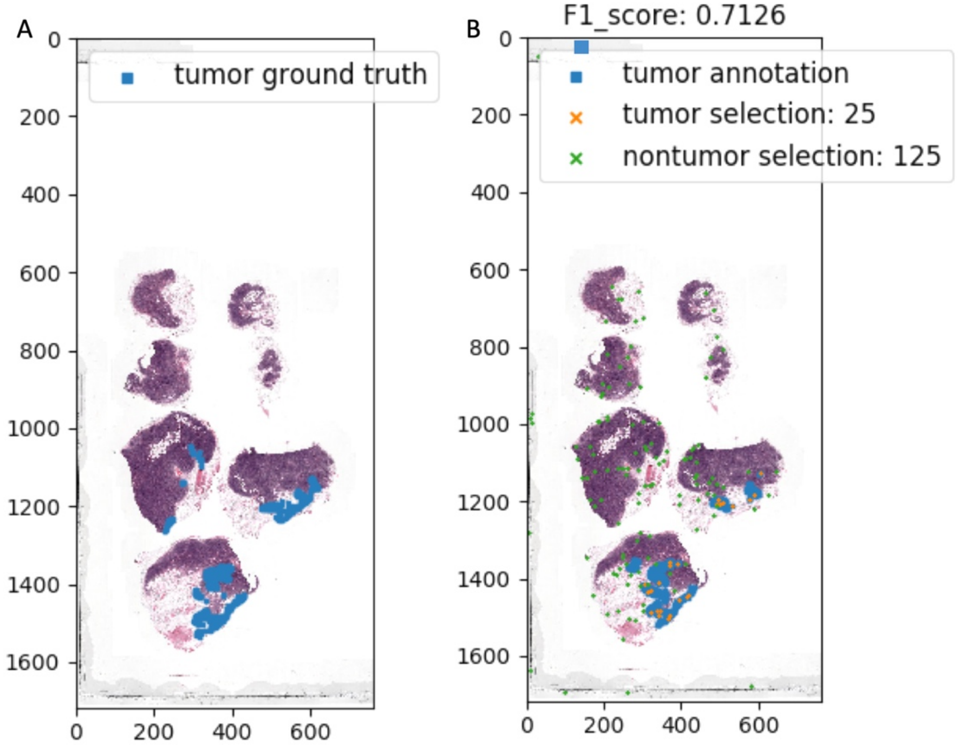


**Fig. S1** *Gold standard tumor masks and annotation mask using developed annotation tool.*

**SUPPLEMENTARY Figure S2**

An example of patch probability being affected by neighboring patches is shown below in Fig. S2 where analysis of the patch (2, 4) (column 2, row 4) in Fig. S2 (a); this patch would have a probability score of 0.75. That is, 0.2 because the patch itself (2, 4) has been determined to be positive by the two- class SVM. 0.1 is added for each top, bottom, left or right positive patch ((1, 4), (2, 5), (3, 4), and (2, 3), respectively), 0.05 is added for the 3 neighboring corner patches (1, 5), (3, 5), and (3, 3). Since (1, 3) is not indicated as positive, 0.05 is not added to the probability score for that patch. Thus, its probability score becomes 0.2 + 4 x 0.1 + 3 x 0.05 = 7.5.
As another example, patch (4, 2) would have a score of 0.5 (0.1 for each of patches (3, 2), (4, 3), (5, 2), and (4, 1) and 0.05 each for corner patches (3, 3) and (5, 3)).

Once the above process has been performed for all test patches, any patches with a probability score of 0.75 or above are considered primary blobs and any patches with a probability score of 0.4 and above are considered supporting areas that draw the convex hall boundaries as in Fig. S2 (b).

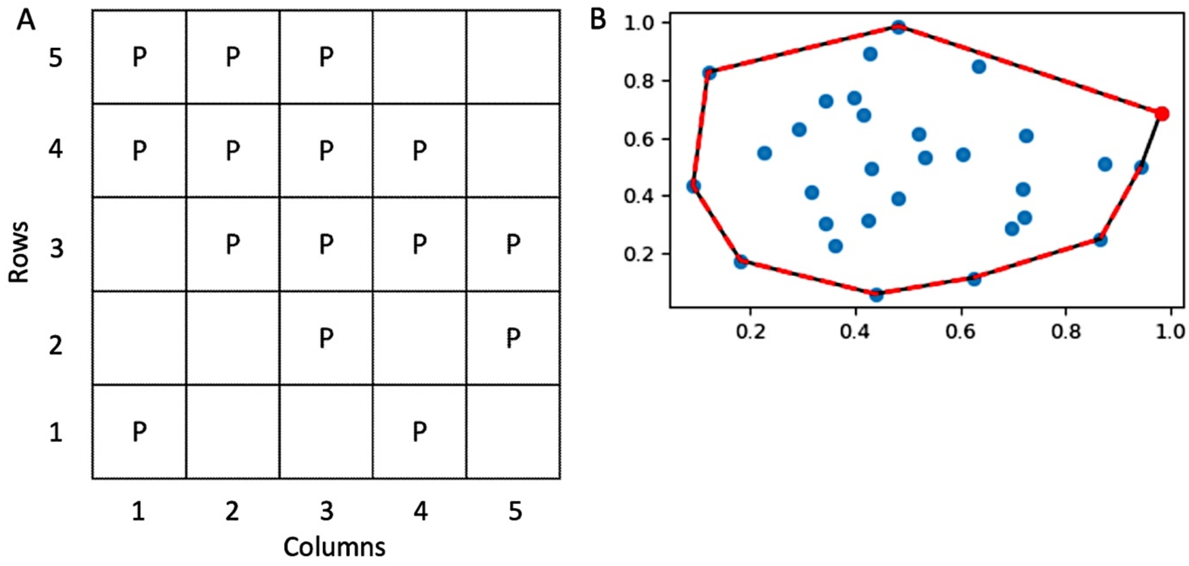


**Fig. S2** *Post-processing using convex hull*. (a) illustrates a representation of positive patches, following analysis by the two-classes SVM, and (b) illustrates a diagram showing a convex hall around a region of interest. [Fig. S2 (b) is from https://docs.scipy.org/doc/scipy/reference/generated/scipy.spatial.ConvexHull.html]
